# First evidence for an aposematic function of a very common color pattern in small insects

**DOI:** 10.1101/2020.07.27.222653

**Authors:** Rebeca Mora-Castro, Marcela Alfaro-Córdoba, Marcela Hernández-Jiménez, Mauricio Fernández Otárola, Michael Méndez-Rivera, Didier Ramírez-Morales, Carlos E. Rodríguez-Rodríguez, Andrés Durán-Rodríguez, Paul E. Hanson

**Affiliations:** Centro de Investigación en Biología Celular y Molecular, University of Costa Rica; Centro de Investigación en Ciencia e Ingeniería de Materiales, University of Costa Rica; Escuela de Biología, University of Costa Rica; Centro de Investigación en Matemática Pura y Aplicada, University of Costa Rica; Escuela de Estadística, University of Costa Rica; Escuela de Física, University of Costa Rica; Centro de Investigaciones en Biodiversidad y Ecología Tropical (CIBET), University of Costa Rica; Centro de Investigación en Contaminación Ambiental (CICA), University of Costa Rica; Protolab. All from the University of Costa Rica

## Abstract

Many small parasitoid wasps have a black-orange-black (BOB) color pattern, which is usually present in both sexes. A likely function of this widespread pattern is aposematic (warning) coloration, but this has never been investigated. To test this hypothesis, we presented spider predators *(Lyssomanes jemineus*), both field-captured and lab-reared individuals, to a species with the BOB pattern and a congeneric all-black species in each of four scelionid genera (*Baryconus, Chromoteleia, Macroteleia* and *Scelio*). Each spider/wasp trial was filmed for 40 minutes under controlled conditions and three behavioral responses (detect, attack, avoid) were recorded in each of 136 trials, never using the same predator and prey more than once. In order to better understand the results obtained, two additional studies were performed. First, the reflection spectrum of the cuticle of the wasp and a theoretical visual sensibility model of the spider were used to calculate a parameter we called “absorption contrast” that allowed us to compare the perception contrast between black and orange in each wasp genus as viewed by the spider. Second, acute toxicity trials with the water flea, *Daphnia magna*, were performed to determine toxicity differences between BOB and non-BOB wasps. By combining the results from the three types of experiments, together with a statistical analysis, we confirmed that BOB color pattern plays an aposematic role.

## Introduction

Many small (<10 mm) parasitoid wasps have a black head, an orange (or reddish orange) mesosoma and a black metasoma. This color pattern has been found in species belonging to 23 families of Hymenoptera (including phytophagous sawflies), and is especially common in neotropical scelionid wasps (Platygastridae; formerly Scelionidae), but is also common in evaniid wasps from diverse biogeographic regions; moreover, this color pattern is usually present in both sexes [1]. In previous research it was found that the spectral blue components of the orange and black color in scelionid wasps are almost identical, suggesting that there is a common compound for the pigments [2], but the identity of the pigment remains unknown.

Conspicuous coloration such as the BOB pattern is often, but not always, indicative of aposematism [3], whereby predators learn to associate particular color patterns with noxious chemical defenses, although this learning process is much more complex than simply developing an aversion to specific types of prey [4]. In some larger insects contrasting black and orange color patterns are known to serve as aposematic (warning) coloration for potential predators, mostly vertebrates [5], and it is possible that the BOB pattern serves as a warning pattern for smaller (invertebrate) predators, although this has not yet been tested.

To test whether the BOB pattern also serves as aposematic coloration for predatory invertebrates, we chose a common jumping spider (Salticidae) as the predator for our experimental trials. Molecular and electrophysiological data suggest that color vision in the principal eyes of most jumping spiders is based on only two types of photosensitive pigments, one sensitive to ultraviolet (UV) light, the other to green light, although a few of them also exhibit filter-based trichromacy [6][7]. These characteristics have allowed salticids to achieve the highest visual acuity thus far measured in any arthropod, with the ability to classify prey into categories, even if they rely solely on vision [8][9][10].

Among the various jumping spiders present in our collecting sites, we chose to use *Lyssomanes jemineus* Peckham & Wheeler, because it was one of the most common species. *Lyssomanes*, like most salticids, are hunting spiders, primarily insectivorous, and behavioral observations suggest a strong role for vision in predation, although visual acuity in *Lyssomanes* may be somewhat lower than that of other salticids, such as *Portia* [11]. *Lyssomanes* species are diurnal foliage dwellers and two hunting behaviors predominate: a sit-wait strategy followed by springing from the underside of the leaf to the upper surface (they usually sit on leaves that are exposed to the sun waiting to ambush prey on the upper surface), and a hunting behavior that consists of exploring both sides of the leaf, actively searching for passing insects [12]. Thus, *Lyssomanes*, like most salticids, can be characterized as a hunting spider that does not use webs, and which executes behavioral responses such as: prey detection by sight, stalking, and attacking by jumping towards the prey [13][14].

The objective of this study was to determine if the BOB color pattern in small parasitoid wasps is aposematic, through toxicity tests and behavioral analysis of invertebrate predators that depend on vision to capture prey. Our hypothesis is that the BOB pattern in scelionids and other small wasps is aposematic, acting as a signal for vision-dependent invertebrate predators. One of our predictions was that when comparing wasp species with the BOB color pattern with others having a black color pattern, the former would show greater toxicity. A second prediction was that in predator choice experiments all-black prey would be preferred over prey with the BOB pattern. Also, experienced predators (collected in the field) would avoid the BOB pattern more than laboratory-bred spiders that were never exposed to it.

## Materials and Methods

### Scelionid collection

The collection site was in the Central Valley of Costa Rica, in a forest patch within the Protected Zone of El Rodeo, which is located near Ciudad Colón, San José (9°52’–9°56’N, 84°14’–84°20’W). The area is composed of secondary forest and a remnant of primary forest (approximately 200 ha). Because the hosts of scelionid wasps (insect eggs) are very difficult to locate, live wasps were obtained via sweeping with insect nets which requires considerable time and effort. Between March 2017 and November 2018 thirty trips were made to the site, and on each visit three people working for about six hours obtained four to six scelionid wasps. The latter were promptly used in trials (see below) carried out in a makeshift laboratory at the same site. Within the wasp genera showing the black-orange-black (BOB) pattern, only some species have this pattern, the other species usually being totally black (Fig 1). The BOB and black wasps were identified by R.M. and P.H. using keys to scelionid genera [15]; voucher specimens were deposited in the Museum of Zoology of the University of Costa Rica. Four genera were collected from this site and used in the experiments: *Baryconus* Foerster, *Chromoteleia* Ashmead, *Macroteleia* Westwood and *Scelio* Latreille. Due to the lack of taxonomic studies for three of the four genera, species level identifications were not carried out (we separated a black morphospecies and a BOB morphospecies in each of the four genera).

**Fig 1.**
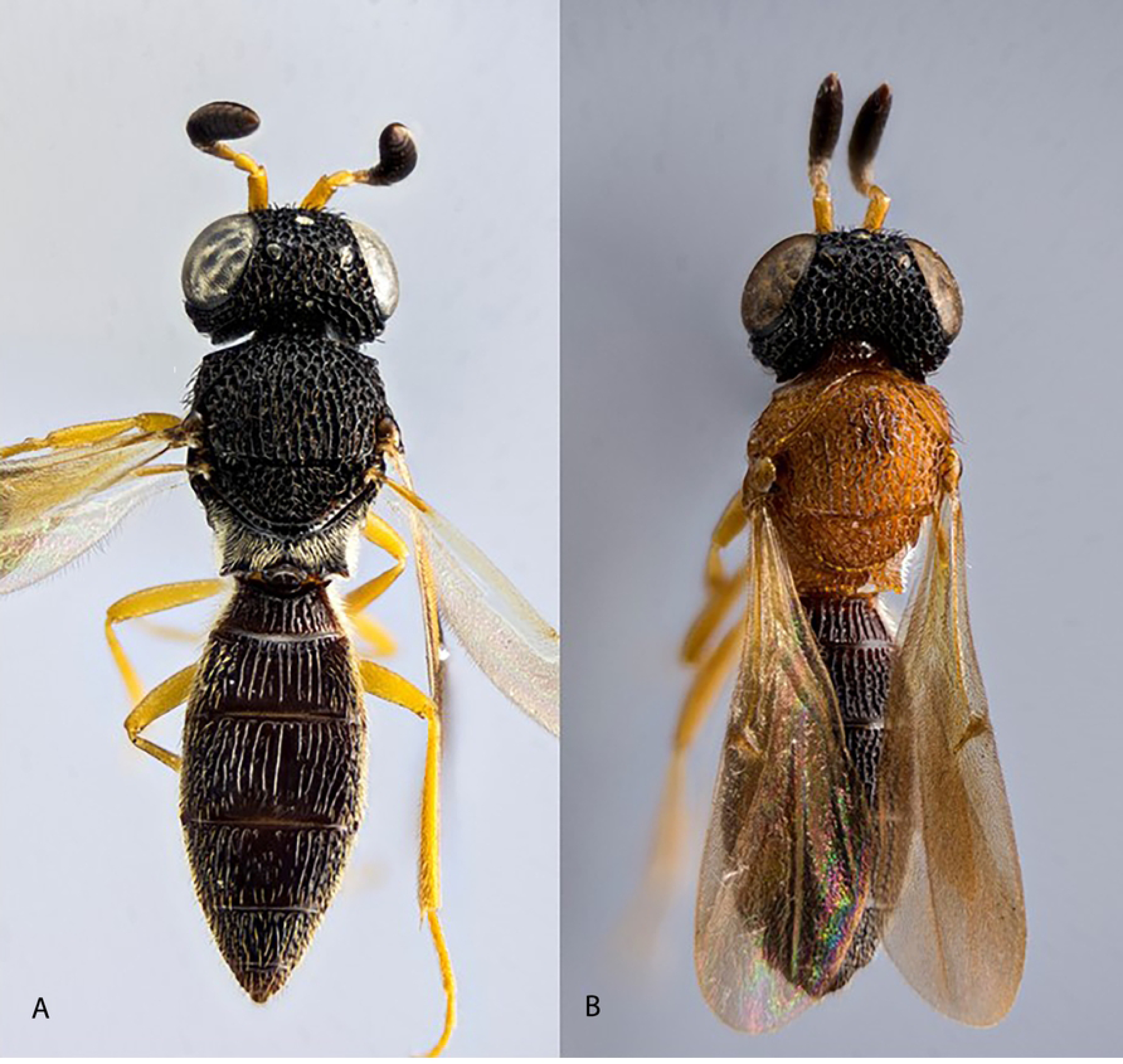
Two scelionid wasp color patterns within the same genus. Color patterns in the genus *Scelio*: black (A), and BOB (B). The focus-stacked macro photograph was obtained with a Reflex 850 camera coupled to a 20x microscope lens.

### Salticid collection

*Lyssomanes jemineus* egg sacs (from which adults were reared in captivity), female adults, and juveniles were collected weekly from January 2017 to January 2019. Collecting was done in Finca 3 of the University of Costa Rica, San Pedro de Montes de Oca, San José, Costa Rica (9°56’07’’N, 84°03’04’’W). This site is located in a disturbed urban environment which exhibits mainly semi-woodland vegetation of native and introduced herbaceous plants [16]. This locality has an average annual precipitation of 1200-1500 mm and an average temperature of approximately 21°C. Spiders and egg sacs were collected from 0 to 1.5 m above ground level. Since many of the plants were quite widely spaced, the transect consisted of 25 to 30 locations where diverse herbaceous plants were haphazardly searched. For the collection of egg sacs weekly samplings of approximately three hours each were carried out from May to September, according to the reported phenology for this species [12]. Each egg sac (usually located on the underside of the leaf) was collected manually without being detached from the leaf and was quickly transferred to the laboratory using 15 ml sterile Falcon tubes with a humid cotton ball. For the collection of adults, the same methodology was applied, but mainly from April to October. Specimens were identified by G.B. Edwards.

### Salticid rearing

Several cage designs were tested, but the most successful (in terms of preventing the escape of spiderlings, optimizing environmental conditions, and facilitating the filming of spider behavior) was one made of acrylic, 14 cm ⨯ 14 cm and 18 cm high, with two lateral circular areas (10 cm diameter) with small holes for aeration covered with an ultra-fine mesh. Including vegetation in the cages helps mitigate the effects of captivity [17], and we found that the best results were obtained by using the same plant species that harbored the egg sac in the field; water was provided in test tubes plugged with cotton. We occasionally observed advanced stage juveniles preying on younger stages, but this behavior was sufficiently uncommon that it was deemed unnecessary to separate specimens individually during the life cycle. However, cannibalism increased as the number of individuals per cage increased and therefore no more than 10 individuals were maintained in a single cage; they were generally separated during the fourth week after hatching, which is when they usually begin to disperse.

A survival rate of 53% was achieved by providing young spiderlings and juveniles (less than 200 days old) with whiteflies, *Bemisia tabaci* Gennadius, and then transitioning to *Drosophila melanogaster* Meigen; older spiders fed with *Drosophila* showed a survival rate of 94%. Whiteflies were reared on young eggplants (*Solanum melongena*) planted in pots inside aluminum cages that were hermetically closed with anti-aphid fabric. Weekly plant watering and room temperature near 24 °C were crucial. In some cases when populations of *B. tabaci* were low we used an undetermined species of *Aleyrodes* found on *Emilia fosbergii, Sonchus oleraceus* and *S. asper*. For *Drosophila* rearing, glass vials with autoclaved *Drosophila* diet (2 bananas, 50 g of oatmeal, 175 ml water and 1.75 ml of propionic acid) were used for rearing at room temperature protected from drafts and direct sunlight.

### Observations of spiders with live prey

Field-collected juvenile and adult spiders, as well as adults reared from egg sacs in the laboratory (for about a year), were observed one at a time in an experimental arena (see below). Adult spiders reared in the laboratory had no prior contact with the wasps. Six types of behavior were recorded: 1) prey detection as indicated by swivelling the cephalothorax; 2) following or stalking prey; 3) crouching, jumping and contacting the prey; 4) piercing and ingesting the prey; 5) prey detection followed by withdrawal from the prey; 6) prey undetected or ignored. In the results the first two were combined as “detect” (although detection was not always followed by stalking), three and four were combined as “attack” (although jumping was rarely followed by ingestion), and five and six were combined as “avoid”.

To facilitate observations of such small specimens we used a white background on the floor, but natural light was allowed to enter the remaining walls of the acrylic box. In order to emulate the natural environment as much as possible, all observations were carried out in a makeshift kiosk embedded in the forest in which the wasps were collected. Behavior was filmed with a Sony RX100 IV camera and was done in FullHD 1080 × 1920 in 60fps. Editing was done in Adobe Premiere, the animations and layout in Adobe After Effects.

In order to minimize potential confounding factors, we used the following standardized conditions for all experiments with spiders.

1. We standardized spider sex (adult females only), age (all juveniles were more than 200 days old) and body size (considered in the statistical analysis).
2. To ensure that the predator was motivated to feed during testing, we fed the spider and then held it without prey for eight days before subjecting it to testing. Eight days also ensured that specimens were not weak, unresponsive or stressed. A faded green body color was sometimes observed in spiders that had fasted for >11 days and appeared to indicate a suboptimal condition.
3. We used each prey and predator for just a single trial.
4. Every time a test spider entered the arena it remained alone for 60 s before introducing the prey, thereby allowing for a short “acclimatization”.
5. Between tests the chamber was washed and cleaned with 80% ethanol, then allowed to dry before beginning the next trial, in order to eliminate any chemical trail of predator or prey since female salticids produce contact and airborne pheromones [18][19][20].
6. Each trial finished when the prey was eaten or after 40 minutes of observation. Preliminary observations prior to the trials revealed that this was a sufficient amount of time, since in the majority of observations all behaviors were usually manifested (even repeatedly) during this interval.

### Observations of spiders with false prey in automated cage

During each trial an adult spider had simultaneous access to two types of lures (painted grains of rice, see explanation below), one painted in the BOB pattern and the other painted all black. The lure moved continuously, one on each lateral wall of the arena, emulating as much as possible a live wasp. All lures were 4 mm in length, similar to that of the scelionid wasps. Data collection was carried out for 40 minutes, similar to the time used for spiders with live prey. We recorded the following behaviors: a) spider detects prey, as indicated by swiveling its cephalothorax; b) prey stalked and/or contacted; c) prey undetected or ignored; d) prey detected and spider withdraws; e) the color pattern (BOB o black) that is detected first. We also measured the body length of each spider and noted whether a silk dragline was produced.

In order to compare spectral characteristics of BOB colors with paints used to color the lures, as well as a possible interaction of these spectral features with the visual system of the jumping spiders, the reflectance spectra of both wasps and paints were considered. With respect to the four scelionid genera included here, we chose to use the previously reported mean reflectance curves for *Baryconus* because this data set showed low dispersion compared with those for other scelionid genera [2]. Six mixtures of different water soluble, solvent-free oil paints (Gamblin®) for the black and orange colors were prepared and tested; the mixtures consisted of combinations between permanent orange, Van Dyke brown, ivory black and Venetian red. Their reflectance spectra were measured using a spectrophotometer (508 PV Craic) coupled to a microscope (Eclipse LV100ND, Nikon). A spectralon standard was used as a diffuse white reference and the experimental setup parameters were the same as in a previous study [2].

Two methods were used for choosing the paint mixture for the lures:

a. Calculating CIELAB color coordinates according to the CIE standard using the reflectance spectra, and then calculating the color differences. In general, color coordinates allow the geometrical representation of colors. In particular, the CIE color spaces are recognized as having important characteristics such as being device-independent and having a perceptual linearity [21]. The CIELAB is a uniform Euclidean space and therefore distances between points can be used to represent approximately the perceived magnitude of color differences between object color stimuli viewed under similar conditions. For the sake of simplicity, in the present work the CIE 1976 L, a, b (CIELAB) color difference definition was adopted, see references in [22][2] for details.
b. Using a color space such as CIELAB implies that colors refer to human visual sensibility, and thus it would be preferable to use information about the visual spectral sensibility of the spider. Nevertheless, to the best of our knowledge, such information has not yet been recorded for the jumping spider used in this work, *L. jemineus*. Preliminary studies of other salticid spiders suggest the presence of photopigments with absorption bands in the UV (ca. 360nm) and the green (ca. 520nm) [23]. Since the spectral information available to us through the use of microspectrophotometry is restricted to the visible wavelengths, we limited our analysis to the use of the nomogram proposed by Govardovskii [24] in order to obtain the normalized absorption spectrum of the αband of A1 type pigment with λmax=520nm. This curve was used to calculate the spectral component of each reflectance curve by multiplying the value of the reflectance curve at each point by the corresponding value of the normalized absorption as described in a previous work [2]. The comparison of the curves obtained in this way for both the reflectance of the BOB pattern of each genus and the reflectance of the paints, allowed us to hypothesize which mixtures would most resemble the wasp cuticle.

### Experimental Arena consisting of an automated cage

The arena for observing spiders with live prey (scelionid wasps) and with false movable prey (lures, explained below) consisted of a closed transparent acrylic box with an internal volume of 1550 cm^3^. The acrylic plates were made in a CNC cutting machine, model Redsail CM1690, with a red laser tube of 100 watts of power. To provide greater structural complexity and additional routes for approaching the prey an acrylic, tree-like structure was placed in the center of the floor [25]. For the experiments with lures we used two types of background: one acrylic box had white-colored walls and the other had black-colored walls.

In the experiments with lures the acrylic box was placed inside a frame made of pressed wood and containing an automated mechanism for moving the lure (Fig 2). The lures were moved by a motor mechanism that allowed for two-dimensional movements. Each lure was connected to a moving part through a nylon thread, generating movement along two axes. This mobile part consists of a magnet on the outside of the acrylic box which in turn moves a magnet on the inside of the box. The control unit has programming functions that allow one to execute different patterns of movement and comprises an interaction interface with the user; for this study, we used only horizontal movements along the side walls of the box (attempting to simulate the movements of live prey that we previously observed to be attractive to the spiders). For the programming of this movement pattern the Arduino platform was used.

**Fig 2.**
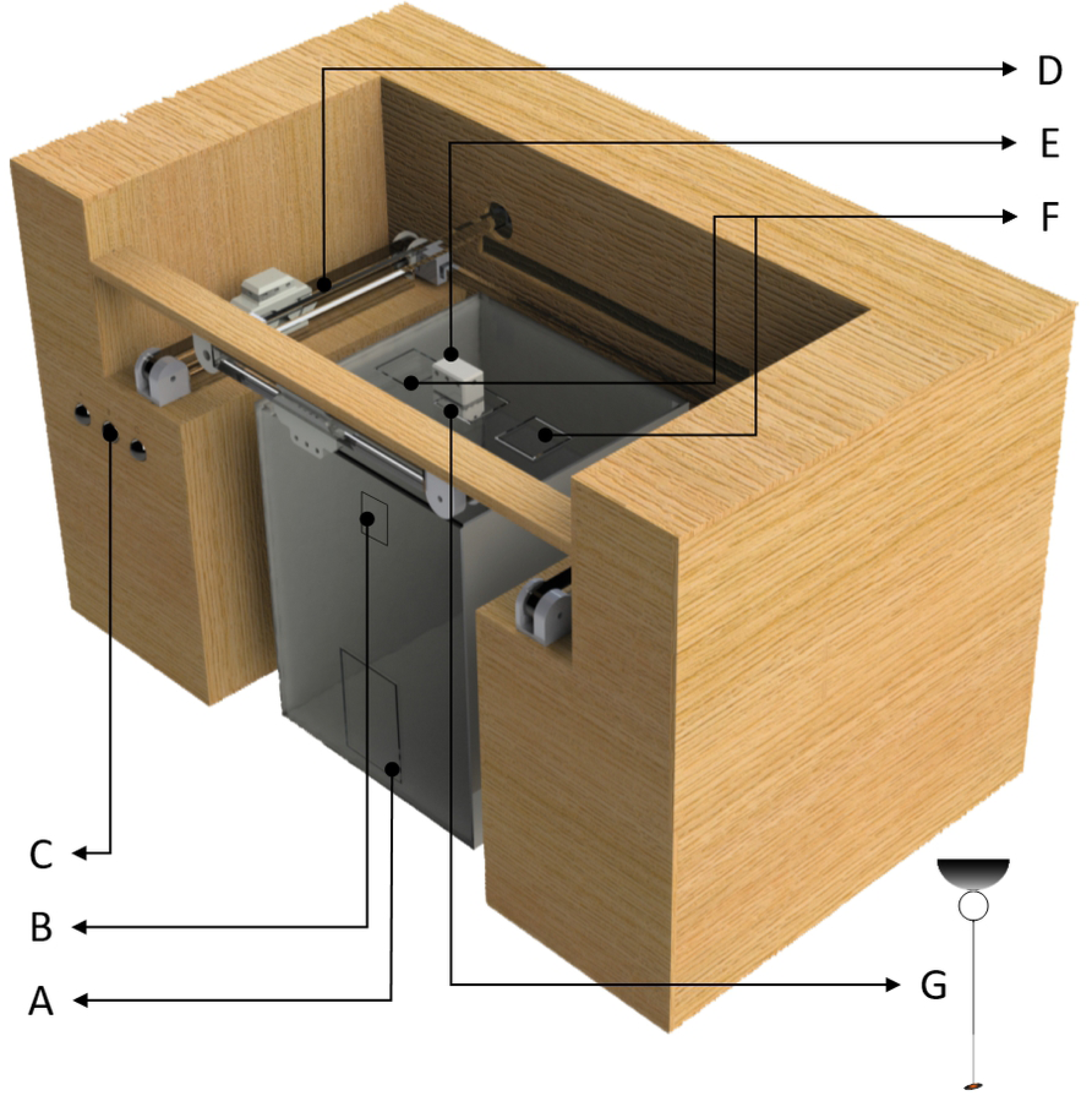
Components of the automated cage. Aperture for introducing the spider (A). Aperture for introducing live prey (B). Buttons controlling the moving lure (C). Motor mechanism consisting of 4 guide blocks, 4 rails and 2 means of movement that uses two pairs of independent rails; each electric motor is connected by a toothed belt to a guide block, which in turn is connected to a busbar that moves the remaining guide block located in parallel; each pair of guide blocks located in parallel is connected to a moving part through a nylon thread, generating movement along an axis (D). Upper magnet (E). Aperture for introducing false prey (lure) (F). Lower magnet from which the lure is suspended by a transparent thread inside the acrylic box (G).

### Green spectral components of the BOB pattern

In order to elucidate whether the spider can distinguish between orange (BOB) and black, and to determine whether the orange color varies between genera, the method described above in item b) was also used to obtain spectral green components of the reflectance spectra of the black and orange color of the BOB pattern in the four wasp genera, as was previously reported [2]. Nevertheless, this description is not complete since it lacks information about the UV visual sensibility.

After calculating the spectral green components, the areas under each curve were obtained numerically. This quantity represents how much of the light reflected by the cuticle of the wasp can be absorbed by the photosensitive pigment of the spider and therefore can be processed by its visual system. According to this criterium, the absolute value of the arithmetic difference between areas should be proportional to a higher or lower capability of detecting a difference between colors. The difference was calculated as the area for the green component of the black color minus the area of the green component of the orange color. Negative values therefore indicate a major absorption of electromagnetic input corresponding to orange, whereas positive values indicate a major contribution to the absorbed light coming from the black color. We refer to these differences as “absorption contrasts”, in the sense that the only assumption we are making about the physiology of the visual system of the spider is the type of photosensitive pigment present and its maximum absorption wavelength (as described above).

### Statistical Analysis

A total of 136 trials with three groups of spiders and live prey were carried out during which the following data were recorded: spider type (juvenile, adult, reared), wasp color (BOB or black), spider behavior (behavioural scale explained before), time when the spider reacted (0-5, 5-10, 10-20, 20-30, 30-40 minutes), the distance between the spider and the prey during each behavior, presence/absence of a silk dragline, and wasp and spider body length. A multinomial logistic regression [26] was fitted to the data, where the behavioral responses were grouped in three general categories (detect, attack and avoid, as explained before) in order to correct for zero or near zero counts in each response category. The remaining variables recorded were included as covariates. Additionally, frequency of actions for each group of spiders were calculated according to time, in order to define a clear timeline of spider behavior in the controlled experiments.

A separate experiment with lures in the automated cage included 30 trials, during which the following variables were recorded: presence/absence of silk dragline, spider size, background experimental arena color (black or white) and color of false prey (BOB vs black) that first attracted the spider.

The response variable in this case was constructed by registering whether *L. jemineus* responded similarly to black and BOB lures coded as 1, and whether *L. jemineus* responded differently to black and BOB lures coded as 2. Contingency tables were calculated, along with a Chi Square independence test for the response versus each of the variables: presence/absence of silk dragline, background color and prey color that first attracted the spider.

Statistical analyses were performed using R (version 3.6.2, R Core Team, 2019). Data formatting and figures were prepared using the Tidyverse packages [27]. Multinomial logistic regression was done using the nnet package [26].

### Wasp extract preparation for toxicity tests

To obtain the wasp extracts some modifications of Arenas research [5] were implemented. Pools of three organisms from the same genus and with the same color were placed in 1.5 ml centrifuge tubes; the initial weight of each pool was taken with a Mettler Toledo XP 205 analytical balance. Extraction was done by adding 0.5 ml of methanol (99.8% purity) to each pool of individuals and subsequent maceration was done with a glass pestle for 5 minutes. After maceration, the tubes were centrifuged in a Minispin plus Eppendorf centrifuge at 14500 rpm for 10 minutes. The supernatant was transferred into a 6 ml glass vial and the pellet was discarded. The methanol from the supernatant was evaporated to dryness with an Organomation brand nitrogen concentrator, model MULTIVAPTM (flow of 7 l/min at 28 °C). For testing, 1400 µl of reconstituted water was added to the concentrate and homogenized using a vortex; this extract was considered as 100 %, then dilutions of 75 %, 50 % and 25 % from the original extract were prepared with reconstituted water to a final volume of 0.5 ml. A negative control of the extract, lacking the wasp pool, was prepared following the same procedure.

### Acute toxicity test with *Daphnia magna*

The toxicity of seven wasp extracts corresponding to four scelionid genera and the two color patterns of interest was measured by an acute toxicity test with the water flea *Daphnia magna* Straus, based on a similar method described by [5]. This organism is frequently used for toxicity tests due to its sensitivity to xenobiotics, wide distribution, short life cycle and ease of culturing in the laboratory. The water fleas were kept in reconstituted water (MgSO_4_, NaHCO_3_, KCl, CaSO_4_ ⨯ 2 H_2_O) [28], at 21 ± 2°C, with a photoperiod of 12:12, and fed with the green algae, *Pseudokirchneriella subcapitata*. The culture medium was changed every week, and after five-weeks adult individuals were discarded and a new culture started from neonates. For the adapted acute toxicity test [29], ten 24-hour neonates were exposed to 0.5 ml of each different dilution (triplicates) of the wasp extract during a 48 hour period in a dark room at a constant temperature of 22 °C. The test end point was mortality (immobilization), which was recorded at 24 h and 48 h. The data were used to calculate the mean lethal concentration (LC50), using the R software, version 3.5.3 (R Development Core Team 2014) and using the “drc” package [30].

### Acute toxicity test with *Vibrio fischeri*

The Microtox test detects inhibition of bioluminescence in the bacterium, *Vibrio fischeri*, using the Microtox® 500 Analyzer. For the analysis, the 2 % Basic Test method recommended by the Microtox® software was employed. Briefly, the lyophilized bacteria (Microtox Acute Reagent) was reconstituted and exposed to four dilutions from the original 100 % wasp extract, prepared with Modern Water Microtox Diluent®. Light emission by the bacteria in contact with the extract dilutions was measured at 0, 5, and 15 minutes; data analysis by the Microtox® software was employed to determine the dilution of the extract that produces a 50% loss of bioluminescence, i.e. the mean effect concentration (EC50).

## Results

### Experiments of spiders with live prey

A total of 136 trials with three groups of spiders (68 field-juveniles, 51 field-adults and 17 captive-adults) were analyzed. For each of the 136 trials consisting of a wasp with a spider, the size of each was measured (Fig 3). Although there is size variation within each group, predator (spiders) and prey (wasps) fall within the same general range, with field-captured adult spiders being the largest.

**Fig 3.**
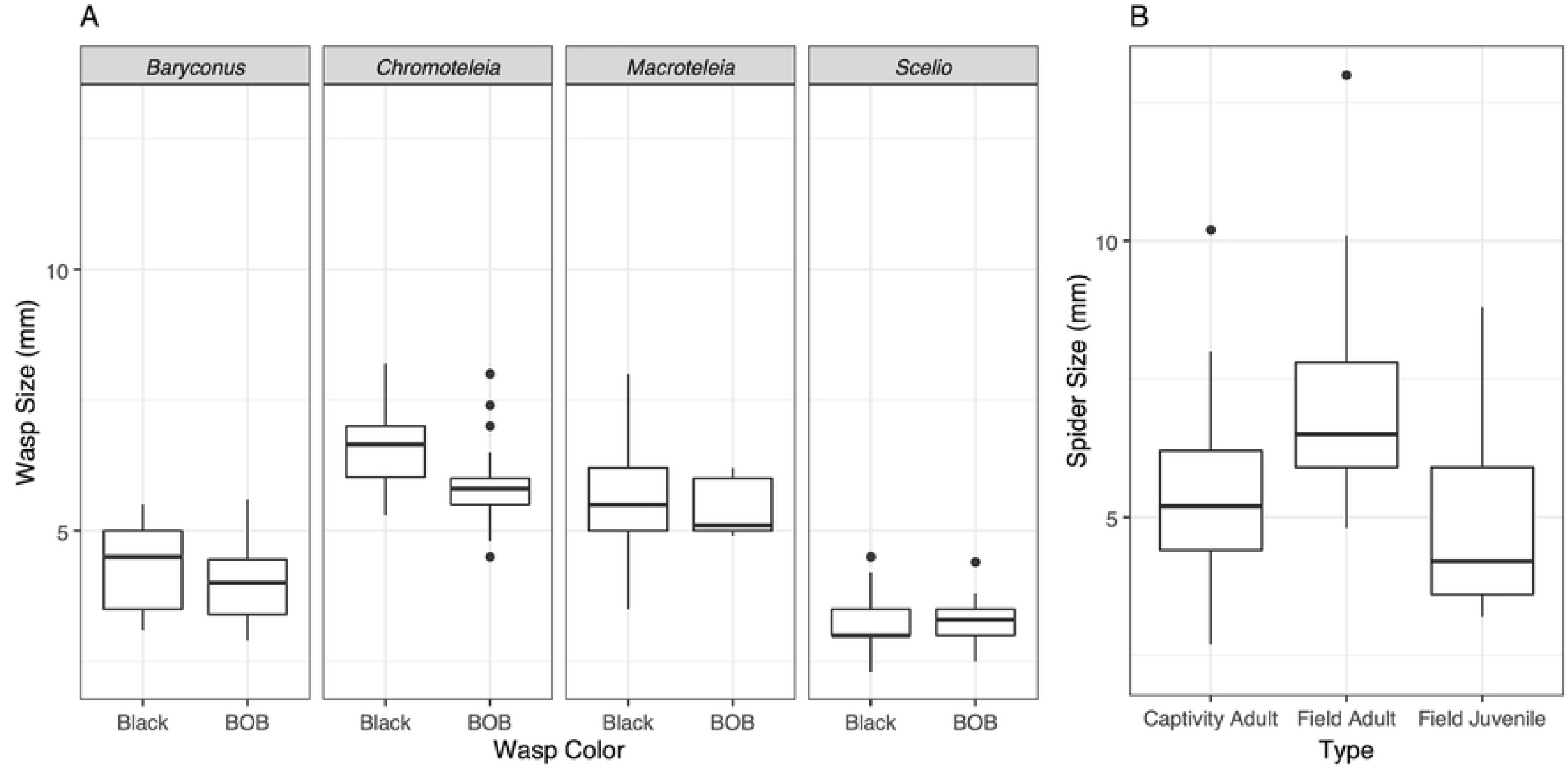
Body length of prey and predator used in this study. (A) wasps according to their genus and color, (B). spiders according to their origin and ontogenetic stage.

For the multinomial logistic regression that was fitted to a simplified response, the most common activity for each spider was classified as “detect”, “attack” or “avoid”. A summary of the data included in the final model is presented in Fig 4. The full model includes the following covariates: wasp color, spider type, wasp size, spider size, wasp genus and presence/absence of silk dragline. This model was compared with simplified versions, with the result being that the model with wasp color and spider type was the model with the best fit, according to the AIC statistic.

**Fig 4.**
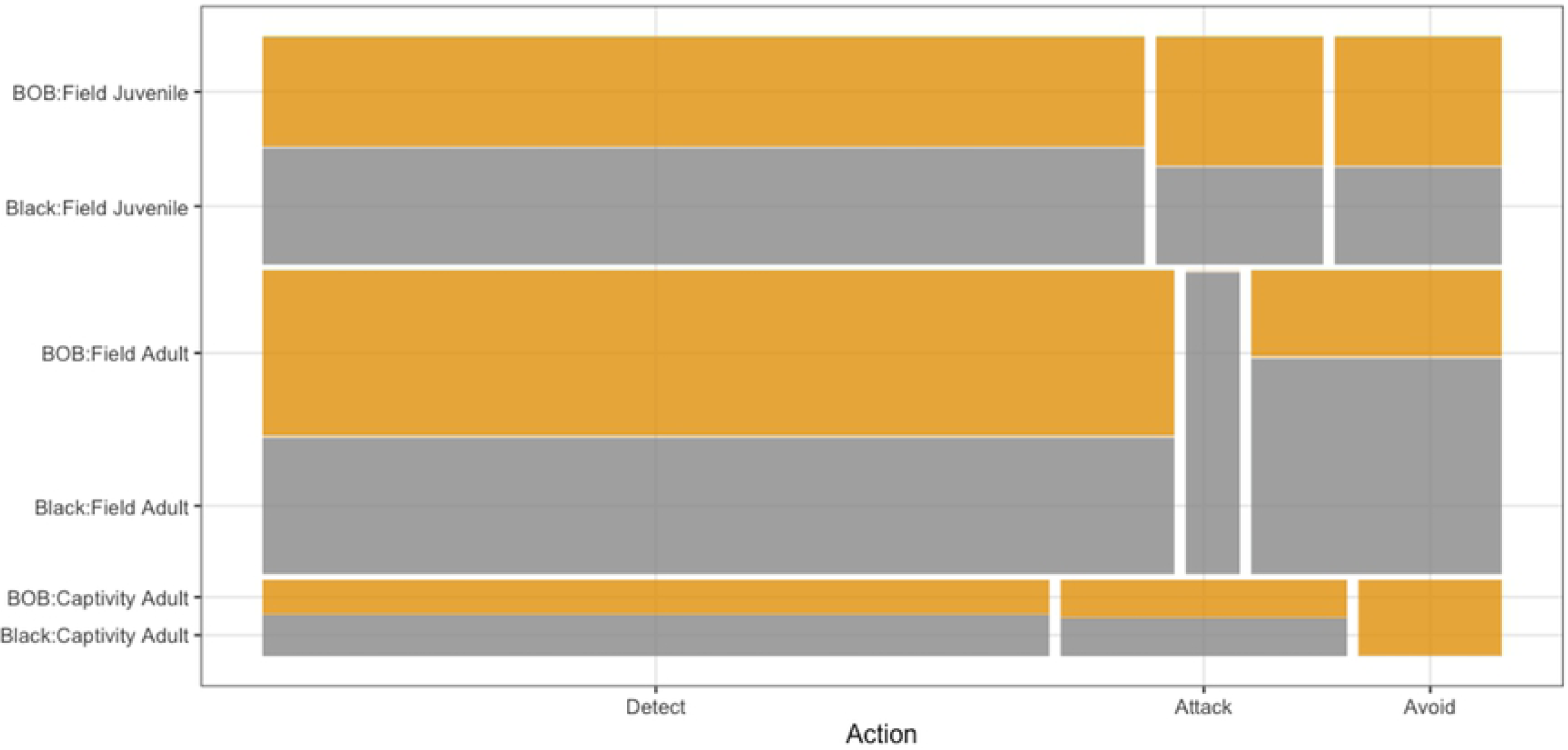
Most common behaviours of *L. jemineus* in trials with different combinations of wasp color (BOB vs black) and spider type. Each bar represents the proportion of spiders per type and wasp color (orange – BOB, gray – black), according to the most common action taken by the spider in a total of 136 trials. The number of spiders differs between groups, which is why the bar for lab-reared (captive) adults is thinner than the ones for field adults and field juveniles.

The aforementioned model suggests that being a field adult is a significant factor for increasing the odds of detecting versus attacking (coefficient = 2.2727, S.E = 0.8922, p-value = 0.0108), and for avoiding versus attacking (coefficient = 2.6374, S.E = 1.1240, p-value = 0.0189). This shows that field adults are more likely to detect or avoid than to attack, compared to adults reared in captivity, which appear to prefer attacking. Similarly, not using silk is associated with an increased chance of detecting versus attacking (p = 0.0160), which makes sense since these spiders use silk only when attacking. Being a field-caught juvenile versus an adult reared in captivity did not make a difference in the most common spider actions. Other factors such as wasp color, wasp size, spider size, and wasp genus did not make a significant difference when included in the full model. It should however be noted that field-caught adults only attacked black wasps, and adults reared in captivity avoided only BOB wasps.

When the behavior of all spiders is analyzed during the 40 min and in the different time slots a greater complexity of behaviors is observed (Fig 5), specifically in that the field adults usually differed from the adults raised in captivity. There is a clear effect of experience, with captive adults showing a lower detection and attack capacity than field spiders (including juveniles). This may be due to the fact that captive spiders were reared on a simple diet and were thus responding to unfamiliar prey. However, there is some evidence for an innate response by spiders in captivity, in avoiding attack responses and aversion to BOB wasps (Fig 5C).

**Fig 5.**
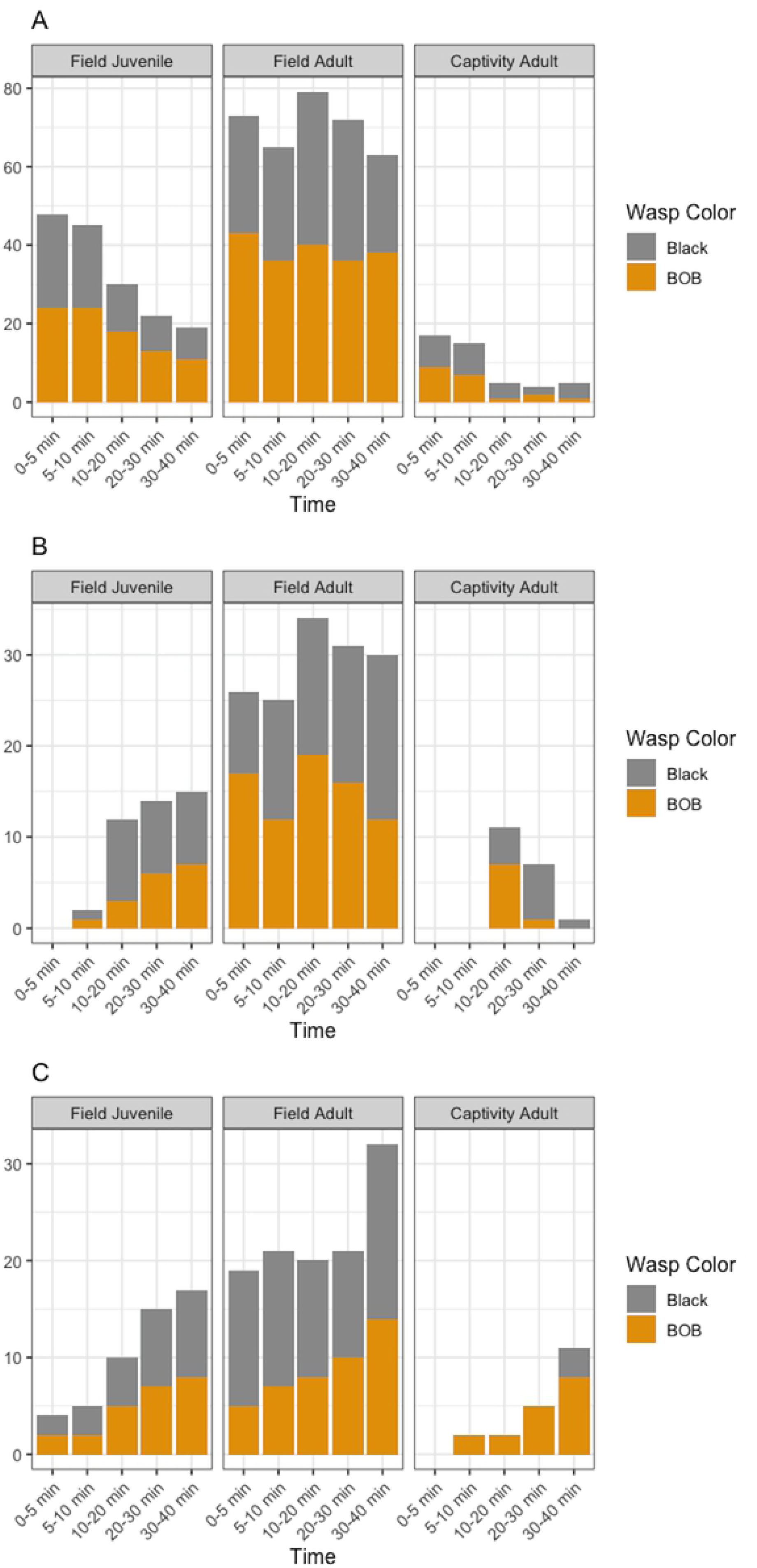
Frequency of behavioral responses experienced by *L. jemineus* when confronted with live wasps. Behaviors are: detection (A), attacking (B), avoiding (C) All bar plots are represented according to behavior per time, spider type and prey color (black and BOB). Time slots without bars indicate that none of the spiders performed that action during that time slot.

Figure 5 summarizes the 954 actions taken by all spiders throughout the timeline of the experiment. There are some time slots without observations: 0-5 mins in field adults, as well as 0-5 mins and 5-10 mins for captive adults in plot B, and 0-5 mins for captive adults in plot C. Thus, the number of spiders for each category changes according to each combination. It can be seen that adults raised in captivity detect/stalk wasps of both color patterns, but during certain time periods their behaviour is directed slightly more toward the black wasps than the BOB wasps (10-20 min and 30-40 min), a behavior not observed in field-adults or field-juveniles (Fig 5A). It is worth noting that adults raised in captivity attack only black wasps during the last time period, unlike the first 30 min during which both colors are attacked, as in field juveniles and adults (Fig 5B). Captive adults avoid wasps with the BOB pattern during most time periods, which differs from the behavior of field-caught spiders, which showed equal avoidance of both colors of wasps (Fig 5C).

### Green spectral components of the BOB pattern

The green spectral components of the black and orange colors of the BOB pattern in the four wasp genera are shown in Fig 6. In general, the spectral components for black and orange colors differ in the position of the maximum wavelength and the height of the peak, indicating different interaction of the reflected light with the photosensitive pigment. The exception is *Baryconus*, for which the spectral components of black and orange colors are similar. In order to provide a quantitative assessment of these differences or similarities, the area under the curve for each spectral component was calculated and the difference (subtraction) between the areas was used as a comparison parameter which was called absorption contrast, as described in the methods. This information, together with the results of the predation experiments, is presented in Fig 7. The four genera were ranked from the highest negative value to the highest positive value of absorption contrast between black and orange spectral components. The information in Fig 7 shows that for the wasps with a BOB color pattern, most of the “attack” events occurred with BOB-colored *Baryconus*, which also has the lowest absorption contrast. On the other hand, the greatest number of “detect” events occurred with BOB-colored *Macroteleia*, which has the highest absorption contrast and the highest contribution of orange, followed by *Chromoteleia* which has the second highest contrast value in absolute numbers, while lower values of black contribution, as in *Scelio* and *Baryconus*, yielded a lower percentage of “detect” events. The greatest number of “avoid” events occurred with BOB-colored *Scelio*, which has the second highest absolute absorption contrast.

**Fig 6.**
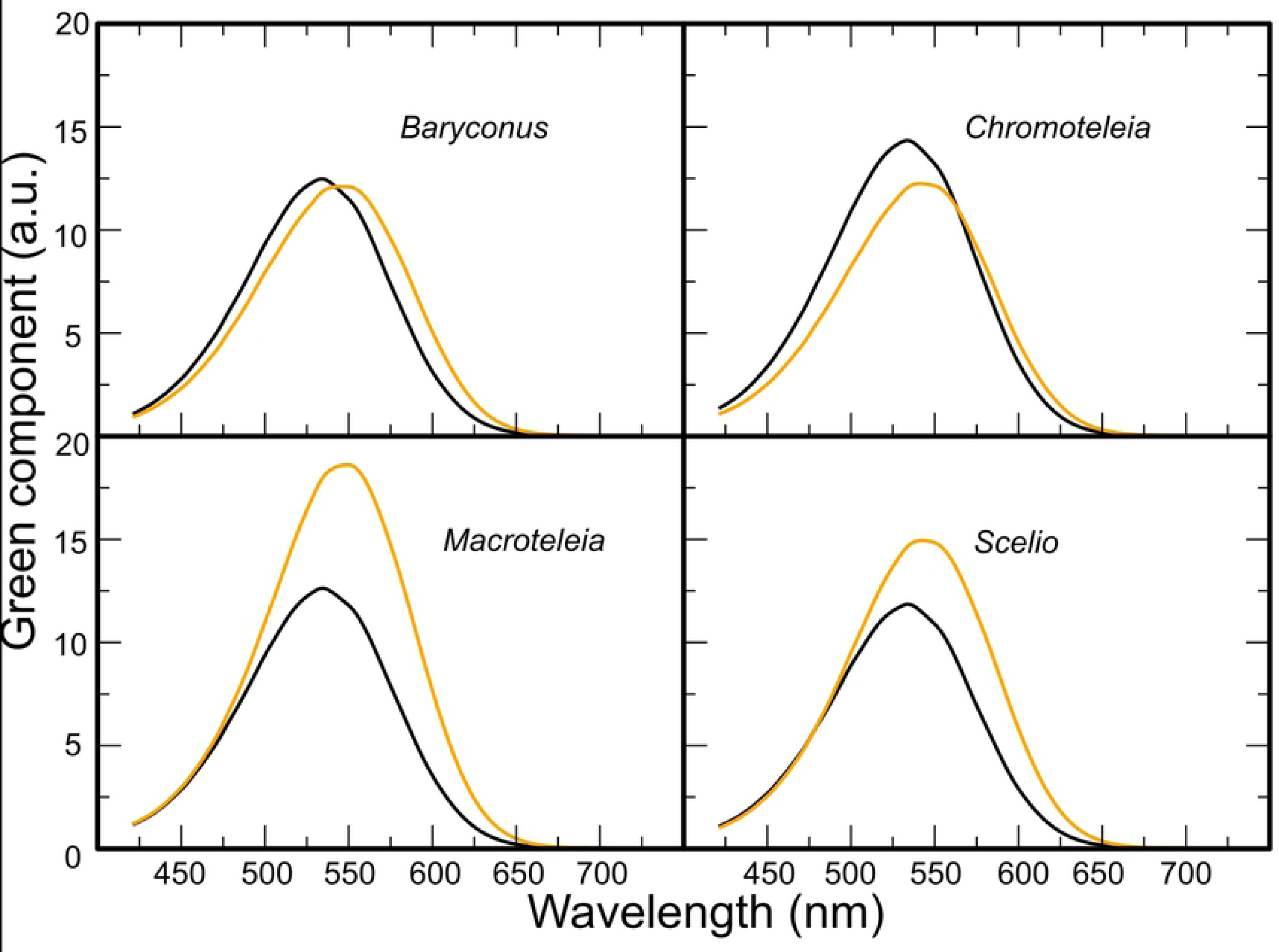
Green component of the reflection spectra of both black and orange colors. Green component, in arbitrary units (a.u.), of both black (black line) and orange (orange line) colors for the BOB color pattern of four genera of scelionid wasps.

**Fig 7.**
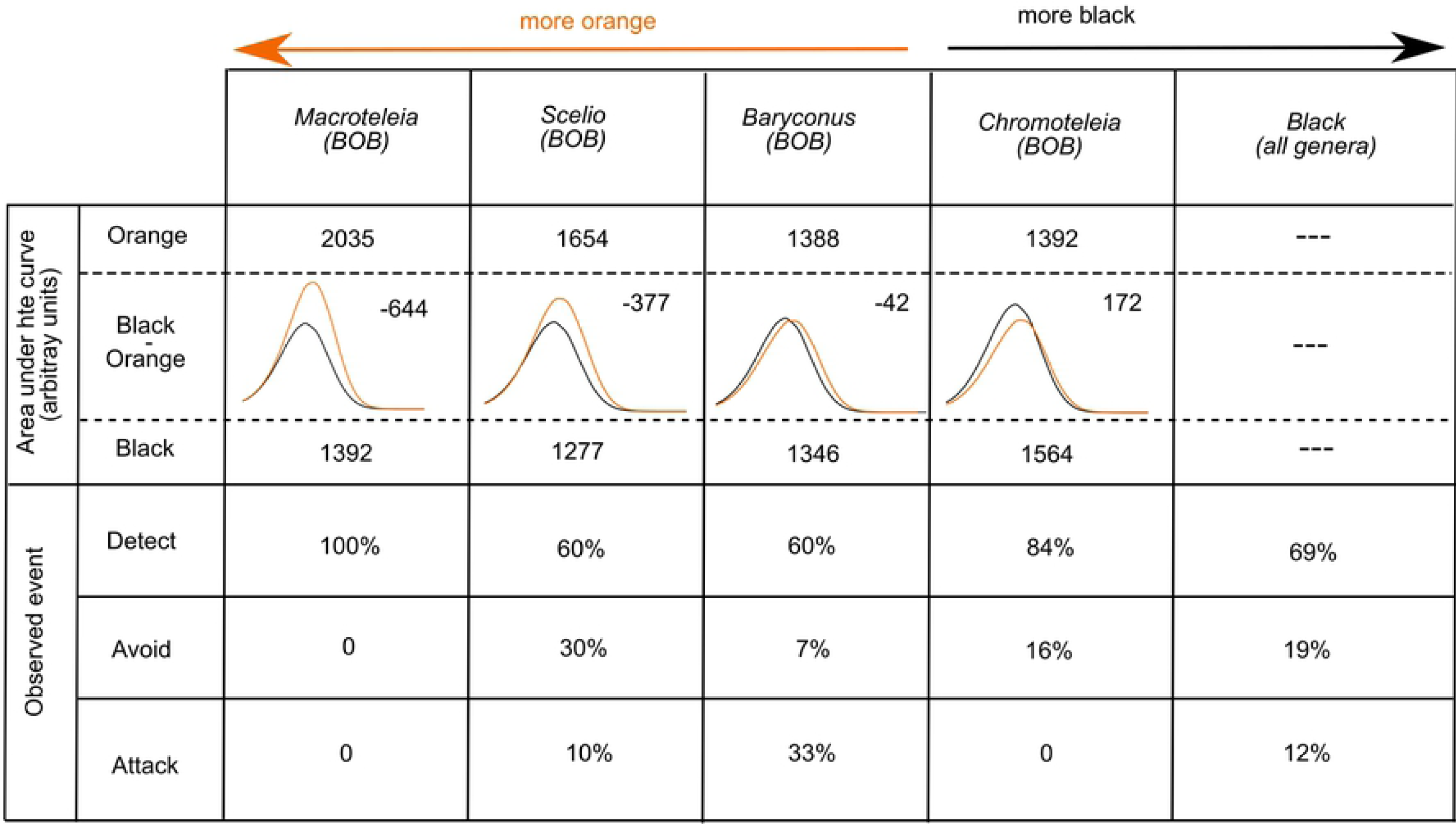
Relationship between the green components of the BOB colors of each scelionid genus and the percentage of behavioral responses experienced by *L. jemineus*. Summary of information about the relationship between the green components of the BOB colors of each scelionid genus and the percentage of behavioral responses experienced by *L. jemineus* observed and grouped in the categories “detect”, “avoid” and “attack”. The BOB color pattern areas under the curve, in arbitrary units, were calculated and compared for the green component of the black and orange colors, respectively (see also Fig 6).

### Experiment of spiders with false prey in automated cage

The reflectance spectra of six blends of black paint and six blends of orange paint were compared with that of the black and orange color, respectively, of BOB-colored *Baryconus* (Fig 8). The best matches in terms of similarity to the black and orange colors of actual wasp cuticle were, respectively, our blend #3 (ivory black + Van Dyke brown) and blend #5 (orange + Venetian red). For black, the difference between wasp cuticle and paint blend was less than 4.5 points, whereas for orange it was 13 points. For reference, 2.3 points is the mean threshold for color differentiation by the human eye [31].

**Fig 8.**
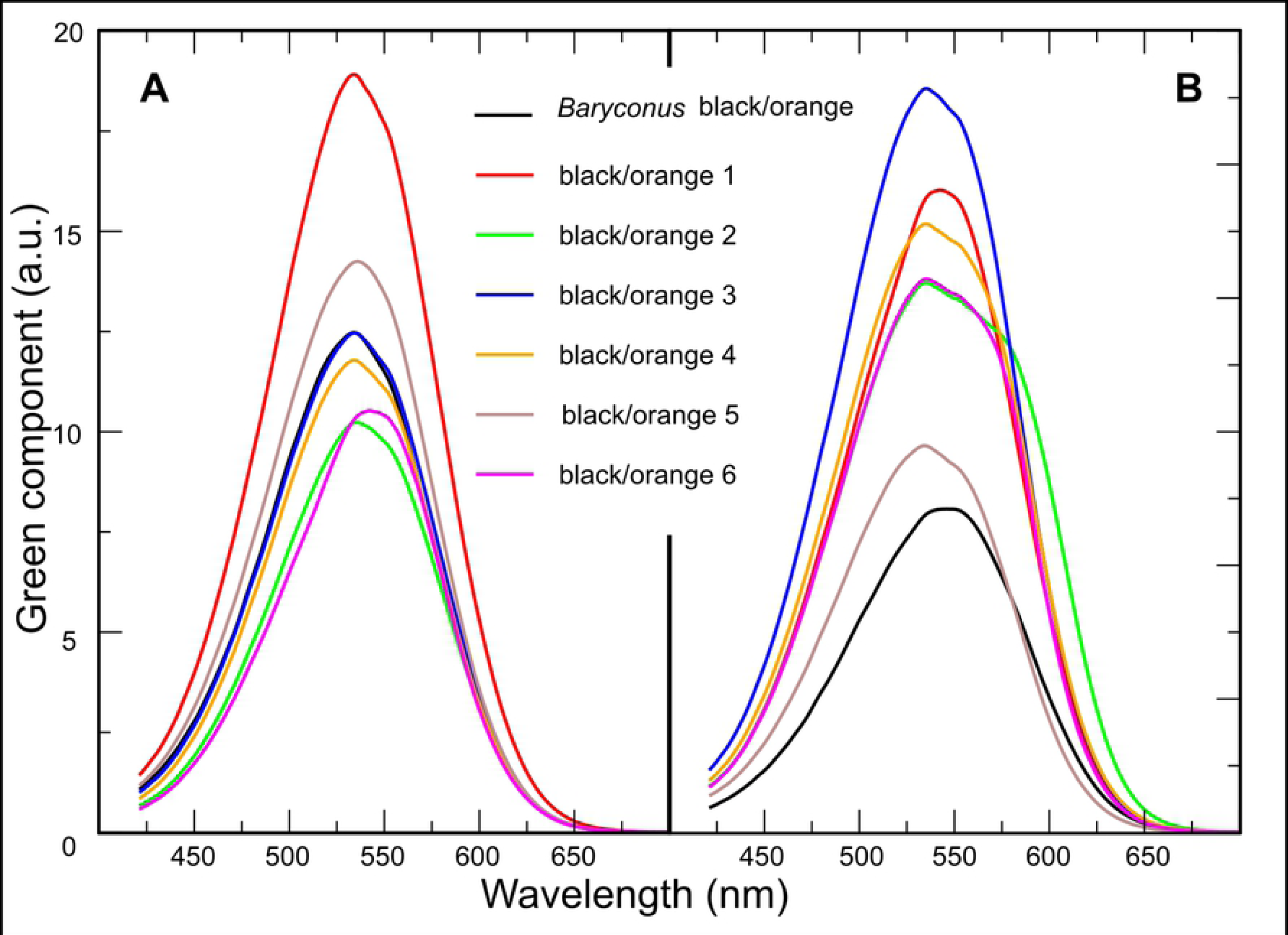
Spectral components, in arbitrary units (a.u.), of the reflectance curves for black-based blends (A) and orange-based blends (B) of paints compared to black and orange colors in *Baryconus*. Note that in (A) the black line for *Baryconus* completely overlaps that for paint 3.

Table 1 presents the results from the experiment with false prey. Each contingency table was tested for independence, and there was no evidence of dependence in any of the cases. In other words, spider behaviors are not associated with background color, the color of lure that was detected first, the use or non-use of silk, or predator size.

**Table 1.**
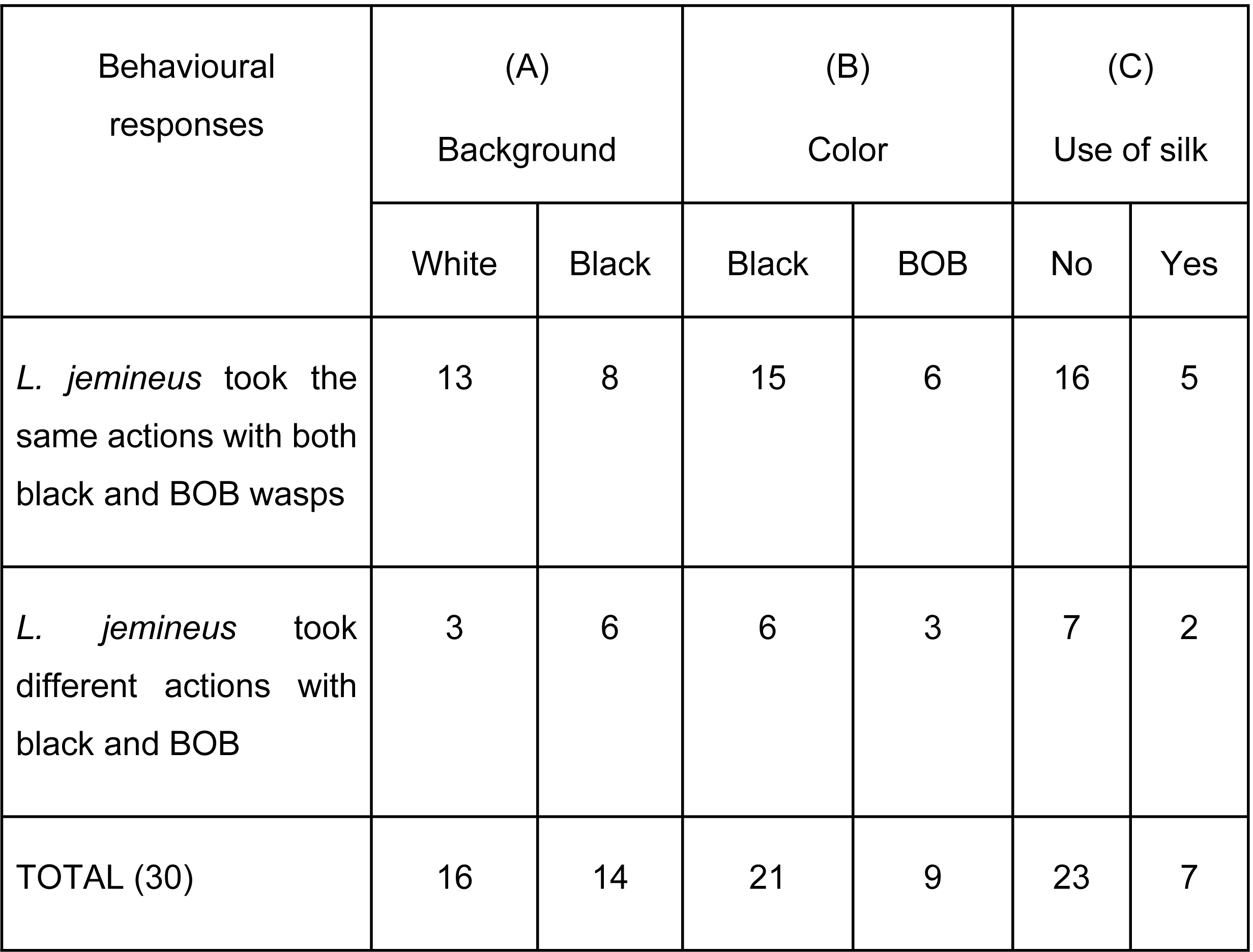
Results from experiment with false prey in three contingency tables. All 30 observations were aggregated according to the response variable (same or different actions) and three variables: (A) background color (white or black), (B) color (black or BOB) that first attracted the spider, (C) use of silk dragline (yes or no).

| Behavioural responses | (A)<br>Background |  | (B)<br>Color |  | (C)<br>Use of silk |  |
| --- | --- | --- | --- | --- | --- | --- |
|  | White | Black | Black | BOB | No | Yes |
| *L. jemineus* took the same actions with both black and BOB wasps | 13 | 8 | 15 | 6 | 16 | 5 |
| *L. jemineus* took different actions with black and BOB | 3 | 6 | 6 | 3 | 7 | 2 |
| TOTAL (30) | 16 | 14 | 21 | 9 | 23 | 7 |

### Acute toxicity tests with *Daphnia magna* and *Vibrio fischeri*

During the exposure period, acute toxicity trials with *D. magna* resulted in a higher mortality (LC50 < 65.2%) of water fleas when exposed to the extracts from the BOB wasps, which employed *Chromoteleia* and *Baryconus* (Table 2). In the case of the black wasps, three out of four extract samples did not show toxic effects (LC50 > 100 %) on *D. magna*; the one sample of black *Macroteleia* that showed toxicity had a lower mortality than with the BOB wasps. No mortality of water fleas was observed with the negative control, thus suggesting that the toxic effects are due to the components of the wasps. In the toxicity tests with *V. fischeri* only black *Macroteleia* and BOB *Chromoteleia* resulted in high bioluminescence inhibition. The results for the two genera have overlapping confidence intervals, so no difference can be inferred; on the contrary, there were no inhibitory effects with BOB *Baryconus* or black *Scelio* (Table 2). Overall, the wasp extracts showed a much higher toxicity for the bacteria bioindicator (*V. fischeri*) than for the arthropods (*D. magna*).

**Table 2.** Toxicity (± I.C.) of extracts of scelionid wasps as determined in acute tests in the bioindicators *Daphnia magna* and *Vibrio fischeri*.

| Wasp genus and color (pools of three specimens) | Total mass (mg) | LC50<sup>a</sup> (%<sup>c</sup>) (*Daphnia magna*) | EC50<sup>b</sup> (%<sup>c</sup>) (*Vibrio fischeri*) |
| --- | --- | --- | --- |
| *Baryconus* (BOB<sup>d</sup>) | 3,41 | 59.6 % (50.2, 68.9) | > 100 % |
| *Chromoteleia* (BOB) | 3,46 | 60.6 % (51.5, 69.7) | 1.7 % (0.9, 3.0) |
| *Chromoteleia* (BOB) | 3,75 | 65.2 % (54.4, 75.9) | 5.2 % (0.8, 32) |
| *Macroteleia* (black<sup>e</sup>) | 1,34 | 85.9 % (73.5, 98.3) | 1.0 % (0.4, 2.0) |
| *Macroteleia* (black) | 1,42 | > 100 % | 2.1 % (1.1, 4.1) |
| *Scelio* (black) | 0,66 | >100 % | > 100 % |
| *Scelio* (black) | 0,86 | >100 % | > 100 % |

<sup>a</sup> LC50: mean lethal concentration
<sup>b</sup> EC50: mean effect concentration
<sup>c</sup> %: Refers to the % of original extract dilution; 100% corresponds to the pure extract.
<sup>d</sup> BOB: black-orange-black color pattern
<sup>e</sup> black color pattern

## DISCUSSION

### Aposematism of the BOB coloration

A previous study showed that the black-orange-black (BOB) color pattern is very widespread among small parasitoid wasps and that it is usually present in both sexes [1], suggesting that this color pattern is not related to sexual behavior. The principal hypothesis regarding the function of this widespread color pattern is aposematism and to the best of our knowledge, the present research is the first to test this hypothesis. Although the results were mixed, at least two of our findings provide evidence that BOB coloration is indeed aposematic. First, although the behavioral responses were quite variable, on five occasions field-captured *Lyssomanes* spiders consumed black wasps (all belonging to the genus *Scelio*), but no spider consumed a wasp with BOB coloration. Second, toxicity, a well-studied and common attribute utilized as a defense by aposematic prey [32], was determined in acute toxicity tests with *Daphnia magna* and showed that wasps with BOB coloration caused greater mortality than did the black wasps. This suggests that BOB wasps contain defensive compounds unlike black wasps, which would explain why only black wasps were consumed.

It was also observed that the wasp extracts were more toxic for the bacteria (*V. fischeri*) than for the crustacean (*D. magna*). Unfortunately, although in the same toxicity range, no clear differences were observed for the extract samples that showed toxicity in the *V. fischeri* assays, due to overlapping confidence intervals. Nonetheless, results from this test should be taken into account when considering the potential presence of toxic compounds in the wasp extracts. A consistent result for both bioindicators is that the black *Scelio* did not show toxicity, and might be considered the least toxic of the four genera. The results for *D. magna* are more relevant for this study since differences between the wasp genera can be observed and because it is an arthropod as is the spider. Moreover, the extracts from the BOB wasps all showed toxicity while just one out of four black wasp extracts did so; moreover, the EC50 values for the BOB wasp extracts were lower, i.e. had higher toxicity.

### Predator responses towards BOB wasps

It should be emphasized that considerable effort is needed to obtain live scelionid wasps; a total of about 540 hours were required to collect the wasps used in our experiments and even with this effort there were insufficient numbers for some of the trials. This limitation was one of the motivations for carrying out trials with lures. Despite considerable care taken in providing lures that closely matched the black and orange colors of the wasps, not subjectively through our visual perception but through spectrophotometric measurements, and an arena in which the movements of the lures were carefully controlled, the results were inconclusive. The likely explanation is that the lures lacked additional visual cues (e.g. legs and eyes) and/or chemical or behavioral cues necessary to elicit a predatory response, which has been shown in other jumping spiders [6]. In future research it would be interesting to use lures simulating additional wasp features, as well as dead scelionid wasps as lures.

In our experiments with predators we used three types of *Lyssomanes* spiders: field-collected adults, field-collected juveniles, and lab-reared (captive) adults. In order to consider warning signals more generally, we integrated background contrast, predator vision, predator life stage, predator and prey sizes, and prior opportunities for learning by the predator [3][5]. Predator origin (being a field adult) was the only significant factor that increased the main probability of detecting vs attacking and avoiding vs attacking. If learning plays a role in avoiding aposematic prey, one would expect the first group (field-collected adults) to show the greatest discrimination between black and BOB wasps, since the use of lab-reared specimens eliminates the role of learned generalization [33], and field-collected juveniles have had less time to learn. The results of our experiments provide partial evidence for this. The only instances of spiders consuming wasps (non-aposematic, black wasps) were of *Lyssomanes* that had been captured in the field. Moreover, field-collected adults were more likely to detect or avoid than to attack, compared to adults reared in captivity which appeared to exhibit a more limited behavioral scale with attack being the most frequent. Thus, adults from the field, which have had more experience, appeared more selective. This is expected, given the complexity of factors and different prey found in natural conditions compared to our artificial breeding environment where the diet consisted of only two types of prey. However, a possible innate effect is evident in the clear aversion of spiders bred in captivity to wasps with the BOB coloration. Furthermore, for spiders from natural conditions the aversion increased over time, and this aversion coincides with the toxicity analyses and the aposematic pattern of the wasps. These results merit further investigation since it is generally assumed that an adverse behavioral response to aposematic coloration is learned, despite the widespread presence of BOB coloration in various taxonomic groups.

### Spectral components of the BOB pattern

Although orange is usually present in aposematic coloration and warning signaling [34][3], our results show that not all BOB specimens were treated similarly by the spider predators. The orange coloration in BOB wasps varies between genera, despite appearing similar to the human eye. It is possible that the difference between the two light absorbing elements (black and orange pigments) within each genus of wasps is important in affecting predator responses. Such differences could favor the increase of conspicuousness and consequently intervene in initial avoidance [35][36]. *Baryconus* had the lowest absorption contrast between black and orange and interestingly, this genus received the most “attack” events. At the other extreme, *Macroteleia* had the highest absorption contrast, as well as the greatest contribution of orange, which perhaps explains why this genus had the greatest number of “detect” events. Therefore, we propose that the contrast difference between black and orange from the point of view of the predator visual capabilities is an important factor to be taken into account in behavioral experiments. This result might also be linked to the results of the toxicity assays, in which the BOB color pattern is associated to an aposematic function.

It has been reported that jumping spiders are less sensitive to the orange end of the spectrum and thus perceive orange cuticles as more achromatic than structures with relatively more reflectance in the green portion of the spectrum [6]. The aforementioned, plus the absence of a red receptor in many spiders, could cause orange objects to be perceived as monochromatically “green” objects, difficult to distinguish chromatically from green foliage [36][37][38]. Our results seem to be consistent with observations that arthropods may be able to perceive long wavelength difference via achromatic or luminance information [39], and that some color patterns may function as multicomponent signals [40][41].

### Conclusion

Most studies of aposematism have used vertebrate predators and larger prey, whereas arthropod predators of smaller prey have generally been neglected [41]. The BOB pattern is extremely widespread in small parasitoid wasps (as well as in a few other insects) and has evidently evolved independently on numerous occasions [1]. To the best of our knowledge, the present study provides the first evidence that this common color pattern has an aposematic function in these small insects.

However, there are still many unanswered questions. The chemical identity of the pigments and the putative noxious compounds are still unknown. How does learning interact with variability in the orange tones and the putative toxins? Finally, it would be instructive to examine the spectral properties of other wasps with this color pattern, as well as the visual capacity and responses of other potential predators.

## Acknowledgements

We give special thanks to Geovanna Rojas Malavasi, Natalia Jiménez Conejo, Paulina Morales Vargas and Anthony Ulate for their support in field data collection, maintenance and rearing of spiders. We also thank Mauricio Valverde Arce for his support in macrophotography and G.B Edwards for his valuable help in the identification of the salticid spiders. We thank our filming crew, Valeria Romero and Soren Pessoa of MANDUCA productions, for their crucial collaboration.

## Author contribution statement

R.M.C Conceptualization, Funding acquisition, Investigation, Methodology, Project administration, Resources, Supervision, Writing – original draft, Writing – review & editing. M.A.C Data curation, Formal analysis, Methodology, Software, Visualization, Writing – original draft, Writing – review & editing. M.H.J Formal analysis, Methodology, Software, Visualization, Writing – original draft, Writing – review & editing. M.F.O and P.H. Methodology, Visualization, Writing – original draft, Writing – review & editing. A.D.R Methodology, Visualization and Writing – original draft M.M.R., D.R.M., C.E.R. Methodology, Visualization and Writing – original draft and Writing – review & editing.

